# One man’s trash is another man’s treasure - the effect of bacteria on phytoplankton-zooplankton interactions in chemostat systems

**DOI:** 10.1101/271791

**Authors:** M Raatz, S Schälicke, M Sieber, A Wacker, U Gaedke

## Abstract

Chemostat experiments are employed to study predator-prey and other trophic interactions, frequently using phytoplankton-zooplankton systems. These experiments often use population dynamics as fingerprints of ecological and evolutionary processes, assuming that the contributions of all major actors to these dynamics are known. However, bacteria are often neglected although they are frequently present. We argue that even without external carbon sources bacteria may affect the experimental outcomes depending on experimental conditions and the physiological traits of bacteria, phytoplankton and zooplankton. Using a static carbon flux model and a dynamic simulation model we predict the minimum and maximum impact of bacteria on phytoplankton-zooplankton population dynamics. Under bacteria-suppressing conditions, we find that the effect of bacteria is indeed negligible and their omission justified. Under bacteria-favouring conditions, however, bacteria may strongly affect average biomasses. Furthermore, the population dynamics may become highly complex resulting in wrong conclusions if bacteria are not considered. Our model results provide suggestions to reduce the bacterial impact experimentally. Next to optimizing experimental conditions (e.g. the dilution rate) the appropriate choice of the zooplankton predator is decisive. Counterintuitively, bacteria have a larger impact if they are not ingested by the predator as high bacterial biomasses and complex population dynamics arise via competition for nutrients with the phytoplankton. Only if the predator is at least partly bacterivorous the impact of bacteria is minimized. Our results help to improve both the design of chemostat experiments and their interpretation and thus advance the study of ecological and evolutionary processes in aquatic food webs.

## Introduction

Highly controllable and easy to handle laboratory experimental approaches are a useful tool to understand complex trophic interactions in natural systems. A prominent representative of these are phytoplankton-zooplankton chemostat experiments which have proven themselves in multiple studies of basic ecological and evolutionary concepts, see e.g. (Novick and Szilard 1950; Fussmann et al. 2000; Yoshida et al. 2003; Becks et al. 2012). Aside from biomass levels, these experiments often focus on patterns in population dynamics, which are fingerprints of interactions between the organisms. While they are undoubtedly able to provide proof-of-concept-like dynamics, chemostat experiments occasionally lack reproducibility, with unexpected experimental runs often not being published, and inference from individual chemostat experiments may be difficult (Bengfort et al. 2017). We hypothesize that bacteria may be one cause of such experimental irregularities.

In numerous chemostat experiments bacteria are an unwanted but often unavoidable and inherent part of the system. While phytoplankton cultures may be run axenically, most zooplankton cultures contain at least parts of the microbiome of the animals (Ishino et al. 2012; Seah et al. 2017). Due to the usually long duration of chemostat experiments also an unintended introduction of bacteria may eventually occur. Phytoplankton exudation and zooplankton excretion drive production of dissolved and particulate organic carbon, providing resources for these bacteria even without an organic carbon source in the growth medium (Vadstein et al. 2003). Bacteria may hamper algal growth by competition for nutrients (Bratbak and Thingstad 1985) and bacterivory can constitute a substantial portion of zooplankton production (Starkweather et al. 1979; Arndt 1993; Ooms-Wilms 1997).

Nevertheless, bacteria are often neglected in chemostat studies. Motivated by earlier experimental investigations (Starkweather et al. 1979; Aoki and Hino 1996; Hino et al. 1997), we challenge this omission and study under which conditions bacteria can substantially influence phytoplankton growth and contribute to zooplankton production, and thereby affect the shape of predator-prey cycles in a typical chemostat experiment.

We include a carefully parametrized microbial loop into a standard phytoplankton-zooplankton chemostat model (Fussmann et al. 2000) (Fig. 1) and study changes in mean biomasses and population dynamics. We analyze how the response of the system to the presence of bacteria depends on experimental conditions, whether the physiological traits of bacteria, phytoplankton and zooplankton favour or suppress bacteria, and how well the zooplankton is able to ingest the bacteria, i.e. its degree of bacterivory. The experimental conditions determine the relative importance of nutrient inflow and losses of nutrients and organisms due to washout on the one hand versus the internal recycling of nutrients and grazing-induced mortality on the other hand. The physiological traits of phytoplankton and zooplankton determine the rate at which organic carbon is produced and the fierceness of the competition for limiting nutrients between algae and bacteria. The degree of bacterivory of the zooplankton determines the grazing pressure on the bacteria. Experimental conditions, physiology and degree of bacterivory thus define the conditions under which bacteria may or may not thrive and impact the system.

**Figure 1.**
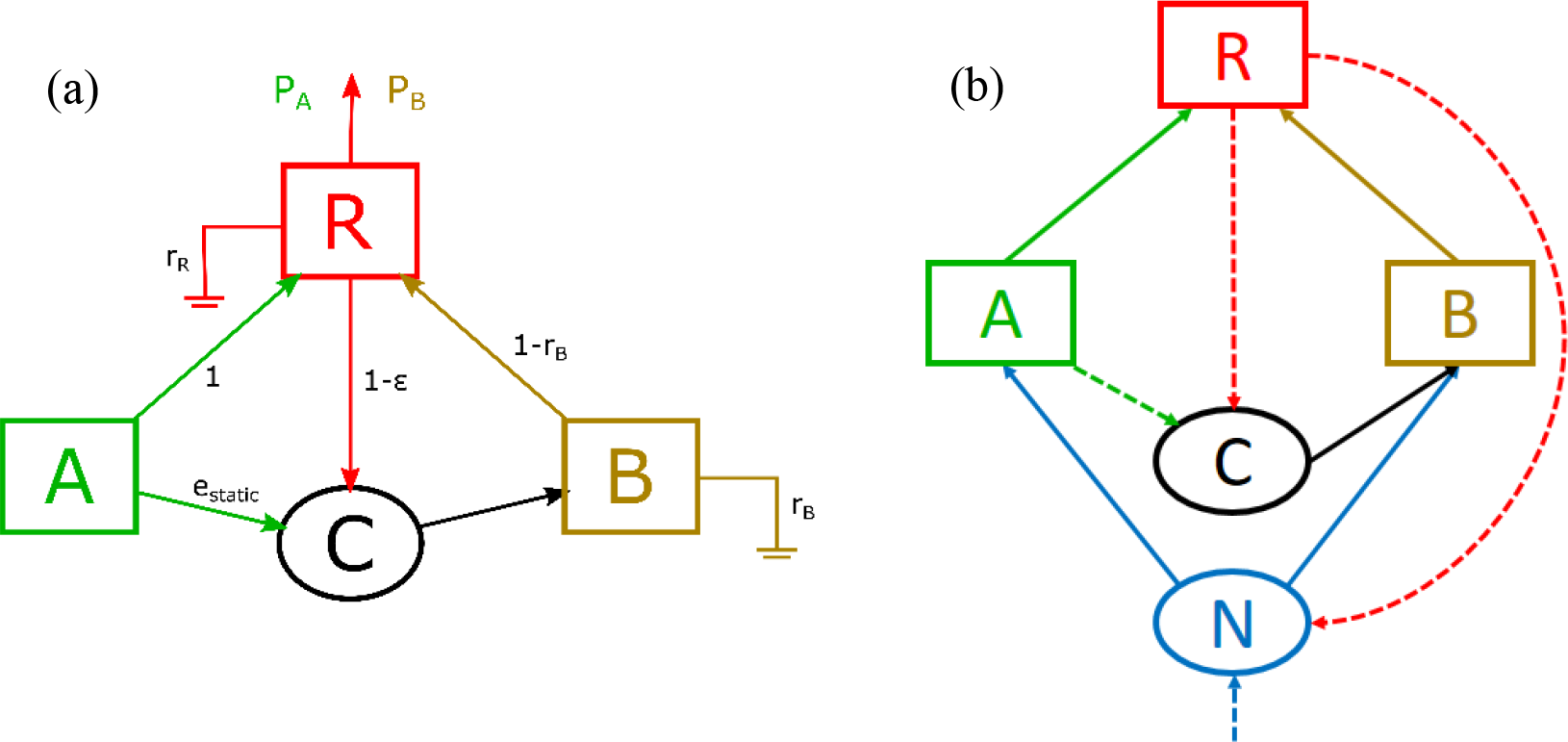
Food web sketches with the limiting resource nitrogen (N), organic carbon pool (C), algae (A), bacteria (B) and rotifers (R). (a) Static carbon flux model with parameters as used in Eqs. 1. (b) Dynamic phytoplankton-zooplankton chemostat model as given by Eqs. 2. Here, solid arrows represent consumption, substance flows are shown in dashed arrows and respiratory fluxes are omitted for clarity.

To sharpen the focus of chemostat experiments on the phytoplankton-zooplankton interaction the question arises how the unwanted but unavoidable effect of bacteria can be minimized. One intriguing strategy to follow could be choosing zooplankton species with a low degree of bacterivory, assuming that non-ingested bacteria would hardly affect the system. Thus, rotifers, which are often less bacterivorous than ciliates (Arndt 1993), may be favoured for phytoplankton-zooplankton chemostat experiments. Instead, we find that the effect of bacteria is low at high degrees of bacterivory. By considering bacteria as inherent actors in phytoplankton-zooplankton experiments we are able to predict conditions when the effect of bacteria is large and provide means to minimize it.

## Materials and Procedures

We employ two models to study the effect of bacteria on phytoplankton-zooplankton interactions. We start with a simple static carbon flux model to obtain general estimates on the dependence of predator production on bacterial production (Fig. 1a). Then, we include more detail and develop a dynamic chemostat model that provides insights into the mean biomasses of all species (Fig. 1b). As a third step, we use this model to study the population dynamics in more detail.

In both models the rate of organic carbon production depends on the physiological parameters for maximal algal exudation *e*_*max*_ and predator excretion (1 - ε), with the predator assimilation efficiency ε. How efficiently this carbon pool can be used by the bacteria is set by their growth efficiency (1 - *r*_*B*_), with the bacterial respiration *r*_*B*_. Bacteria are suppressed by low carbon supply, which is achieved at low algal exudation and low predator excretion, and inefficient use of that carbon by the bacteria. Bacteria are favoured by high carbon supply, i.e. at high algal exudation, predator excretion and bacterial growth efficiency. Using the lower and upper end of the broad ranges of published values for these parameters (Tab. 1) we construct a minimum and a maximum impact scenario of conditions suppressing or favouring bacteria, respectively.

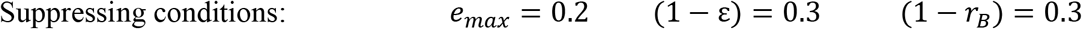

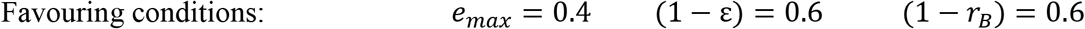

**Table 1.** Parameter values and their biological meaning. Ranges are given for parameters that are varied within this study. Parameter values are either derived from own, earlier chemostat runs, set to reasonable numbers, or taken from literature.

| Parameter | Biological meaning | Value | Reference |
| --- | --- | --- | --- |
| $\delta$ | Chemostat dilution rate | 0.07 .. 0.7 d<sup>-1</sup> | varied within standard experimental ranges (Fussmann et al. 2000) |
| $N_I$ | Inflow resource concentration | 160 $\mu\text{molN L}^{-1}$ | set to standard experimental conditions |
| $\omega_{A,N}$ | $N$ content in an algal cell | $4.6 \times 10^{-8} \mu\text{molN}$ | estimated from own data |
| $\omega_{B,N}$ | $N$ content in a bacterial cell | $8.8 \times 10^{-10} \mu\text{molN}$ | from $\omega_{B,C}$ with C:N = 5.65 (Vrede et al. 2002) |
| $\omega_{R,N}$ | $N$ content in a rotifer | $1.2 \times 10^{-3} \mu\text{molN}$ | estimated from own data |
| $\omega_{A,C}$ | $C$ content in an algal cell | $5 \times 10^{-7} \mu\text{molC} \triangleq 6 \text{ pgC}$ | estimated from own data |
| $\omega_{B,C}$ | $C$ content in a bacterial cell | $5 \times 10^{-9} \mu\text{molC} \triangleq 60 \text{ fgC}$ | set to 1/100 of $\omega_{B,C}$ in agreement with Vrede et al. (2002) |
| $\omega_{R,C}$ | $C$ content in a rotifer | $6.7 \times 10^{-3} \mu\text{molC} \triangleq 80 \text{ ngC}$ | from $\omega_{R,N}$ with C:N = 5.6 (Jensen et al. 2006), in agreement with Dumont et al. (1975) and Rothhaupt (1990) |
| $e_{max}$ | Maximum carbon exudation of algae | 0.2 .. 0.4 | varied (Baines and Pace 1991; Tittel et al. 2012) |
| $e_{min}$ | Minimum carbon exudation of algae | 0.1 | varied (Baines and Pace 1991; Tittel et al. 2012) |
| $r_B$ | Carbon respiration by bacteria | 0.4 .. 0.7 | varied (del Giorgio and Cole 1998) |
| $r_R$ | Carbon respiration by rotifers | 0.5 | Humphreys 1979 |
| $\varepsilon$ | Carbon assimilation efficiency of rotifers | 0.4 .. 0.7 | varied (Straile 1997, Verschoor et al. 2007) |
| $\beta_A$ | Maximum algal growth rate | 1.9 d<sup>-1</sup> | estimated from own data |
| $\beta_B$ | Maximum bacterial growth rate | 1 d<sup>-1</sup> | (Morris and Lewis 1992) |
| $G$ | Rotifer maximum ingestion rate | $3.6 \text{ d}^{-1} \triangleq 288 \text{ ngC d}^{-1}$ | (Rothhaupt 1990a) |
| $H_A$ | Algal half-saturation | $49 \mu\text{molN L}^{-1}$ | estimated from own data |
| $H_{B,N}$ | Bacterial half-saturation for nitrogen | $4.9 \mu\text{molN L}^{-1}$ | set to 1/10 of the algal half-saturation |
| $H_{B,C}$ | Bacterial half-saturation for carbon | $0.83 \mu\text{molC L}^{-1}$ | (Tittel et al. 2012) |
| $H_R$ | Rotifer half-saturation | $195 \mu\text{molC L}^{-1} \triangleq 2.34 \text{ mgC L}^{-1}$ | (Rothhaupt 1990b; Fussmann et al. 2000), verified with own data |
| $p_B$ | Degree of bacterivory of the rotifer | 0.01 .. 1 | varied |

### Static carbon flux model

A first and easily obtained estimate of the effect of bacteria for the extreme cases of suppressing and favouring physiological conditions for bacterial growth in a chemostat may result from a static model of the carbon fluxes between algae, bacteria and rotifers. Thus, we employ a model similar to those presented by Anderson and Ducklow 2001 and Gaedke et al. 2002 to estimate the amount of predator production based on bacterial production for different physiological parameters, i.e. bacteria-suppressing or -favouring conditions (Fig. 1a). Assuming steady state, algal production is ingested and converted into predator excretion (1 − *ε*), respiration (*r*_*R*_) and production. Algal exudation, which is assumed to be proportional to algal production, and predator excretion supply the organic carbon pool. Bacteria consume the carbon pool and may be ingested by the predator. The loop of carbon excretion from feeding on bacteria and subsequent recycling of this carbon by bacteria is resolved by a geometric series. The predator production per unit algal production from algae (*P*_*A*_) and bacteria (*P*_*B*_), respectively, thus becomes

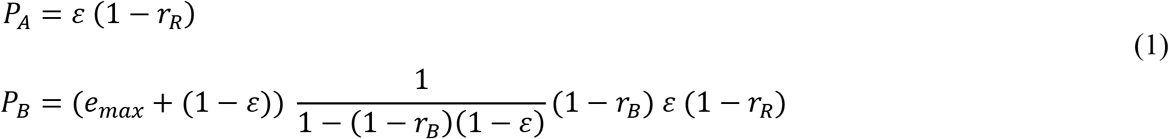

The dependence of predator production on *e*_*max*_, ε and *r*_*B*_ is shown in Suppl. Fig. 1. This model predicts that under suppressing conditions 14% of the total predator production originate from ingesting bacteria. This results in an increase of total predator production by 16%. Under favouring conditions bacteria constitute 48% of predator production which almost doubles with an increase by 94%. From this strongly simplified model we already see that bacteria may have a large effect under certain physiological conditions. As this model conveys no information on actual biomasses or population dynamics we consider below a mechanistic
differential equations model (Eqs. 1) for a full picture of the effects of bacteria on phytoplankton-zooplankton interactions.

### Dynamic phytoplankton-zooplankton model with organic carbon pool and bacteria

Using a well-established model presented by Fussmann et al. 2000 and Yoshida et al. 2003, we describe the predator-prey interaction of the rotifer *Brachionus calyciflorus* (*R*, Ind./L) feeding on the unicellular green algae *Monoraphidium minutum* (*A*, cells/L) in a chemostat (Fig. 1b). We simplify the original model slightly by assuming that every rotifer individual is fertile, i.e. we neglect the short periods of juvenile growth and senescence, but extend it by adding a pool of organic carbon *C* (μmol L^−1^) and bacteria *B* (cells/L). Nitrogen *N* (μmol L^−1^) is the limiting resource for algal growth. Bacterial growth is assumed to be multiplicatively co-limited by nitrogen and carbon. The full model reads

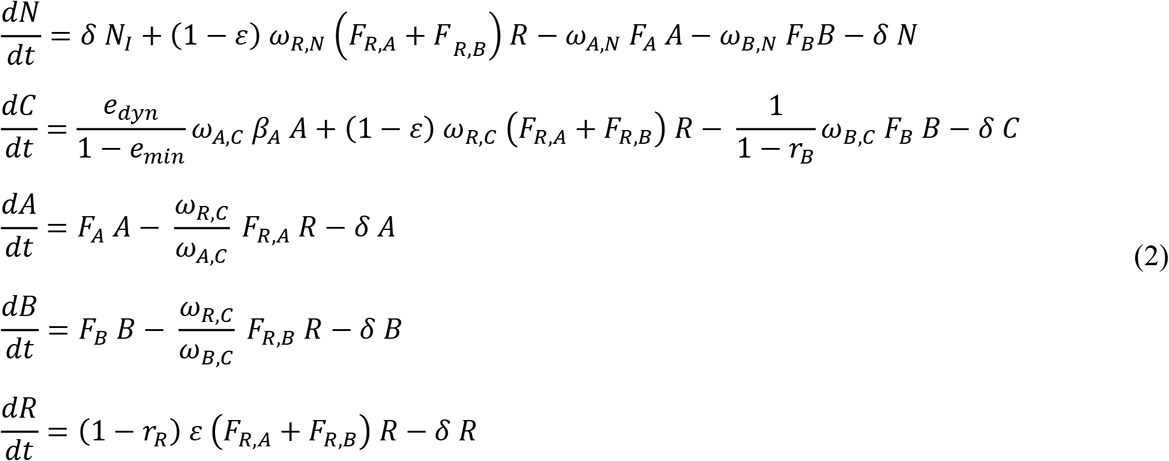

All parameter values are listed in Tab. 1 along with their biological meaning. We will now describe the terms of the model in the order as they appear in Eqs. 2.

The nitrogen pool in the chemostat is supplied by the inflow of fresh medium, which is given by the product of chemostat dilution rate δ and nitrogen concentration in the medium *N*_*I*_, and the excretion of the rotifers from feeding on algae and bacteria. Algal and bacterial growth, at per capita rates *F*_*i*_ (Eqs. 3), and wash-out reduce the nitrogen pool.

The carbon pool is maintained through dynamic, nutrient dependent exudation by algae (*e*_*dyn*_, for details see Appendix 2 and Suppl. Fig. 2) and excretion by rotifers. It is diminished by bacterial consumption and wash-out. The interactions of species *i* with the carbon and nutrient pools are scaled by the respective carbon (ω_*i,C*_) and nitrogen (ω_*i,N*_) content of an individual. Algae and bacteria grow at per capita growth rates [d^−1^] of

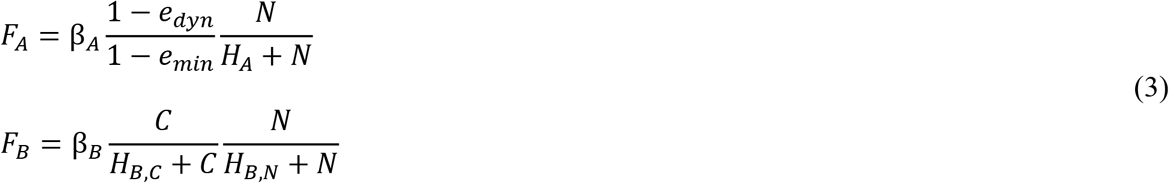
where β_*i*_ is the maximum growth rate of species *i*, *e*_*min*_ is the minimum exudation and *H*_*i*_ is the half saturation constant. Algal and bacterial densities are reduced by rotifer grazing and washout. The rotifer per capita grazing rates on algae and bacteria [d^−1^] follow a multi-species Holling Type-II shape.

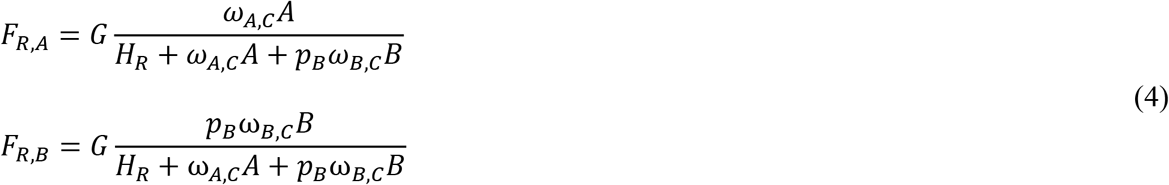

Here, *G* is the maximum grazing rate of a rotifer and *H*_*R*_ is the half saturation constant scaled to carbon. The bacteria are potentially less edible than the algae, depending on the degree of bacterivory of the rotifers *p*_*B*_ which provides the part of the bacterial population that is accessible to the predator. Effectively, this scales up the half-saturation constant of the predator for feeding on bacteria. Grazing is converted into bacterial or algal losses by the ratio of carbon contents per individual.

The rotifers assimilate only a part of the ingested food. What is not assimilated is excreted and enters the carbon pool. The assimilates are further reduced by respiration, the remainders are used for production of new rotifer biomass. The only loss-term of rotifers is wash-out.

### Numerical simulations and determination of dynamics

To achieve a broad picture of the effects of bacteria, we examined the parameter space spanned by the dilution rate of the chemostat and the degree of bacterivory of the predator, thereby considering the two scenarios suppressing or favouring bacteria. The dilution rate is an important parameter for the performance of the individual species as it determines the rate of nutrient input and the loss rates of all species. The degree of bacterivory is important as it shapes the interspecific interactions via the apparent competition between algae and bacteria mediated by the predator. A third important system parameter is the nutrient inflow concentration, which we included in our analysis at an intermediate dilution rate for favouring conditions.

The system of ordinary differential equations Eqs. 1 was integrated with the *odeint* package from the Scipy library (Jones et al. 2001) in Python (version 3.5).

The presence of bacteria in an algae-rotifer system may have two ecologically important effects, first on the mean biomasses, which can directly be obtained from the model outputs, and second on the population dynamics. To distinguish between steady state, regular cycles and irregular dynamics, local peaks in the normalized autocorrelation function (nACF) of the algal density were detected using the *argrelmax* algorithm from Scipy. A time series was classified as being at steady state if no peaks with prominence above the accuracy of the solver were detected. If the first peak of the nACF was above 0.95 the dynamics were classified as regular and the number of algal maxima within one such repetitive unit was obtained. If all peaks of the nACF were below 0.95 the dynamics show no clear repetitive pattern and thus were classified as irregular.

## Assessment

### Effect of bacteria on mean biomasses

Using the mechanistic model, we compare the effect of bacteria in chemostat experiments under suppressing and favouring conditions for large ranges of the chemostat dilution rate and the degree of bacterivory of the predator. These two key parameters, which govern the fluxes in the system, may strongly affect the mean biomasses of all species (Fig. 2). Comparing the two extreme cases of bacteria-suppressing versus bacteria-favouring conditions shows that under suppressing conditions bacterial biomass remains mostly negligible and algal and rotifer biomass is thus independent of the degree of bacterivory by the predator (Fig. 2a). With little bacteria present algal mean biomass increases and predator biomass decreases as the dilution rate increases. Only at very low degrees of bacterivory and high dilution rates the bacteria can achieve non-negligible biomasses, which is reflected by a slightly lower algal biomass in this parameter region.

**Figure 2.**
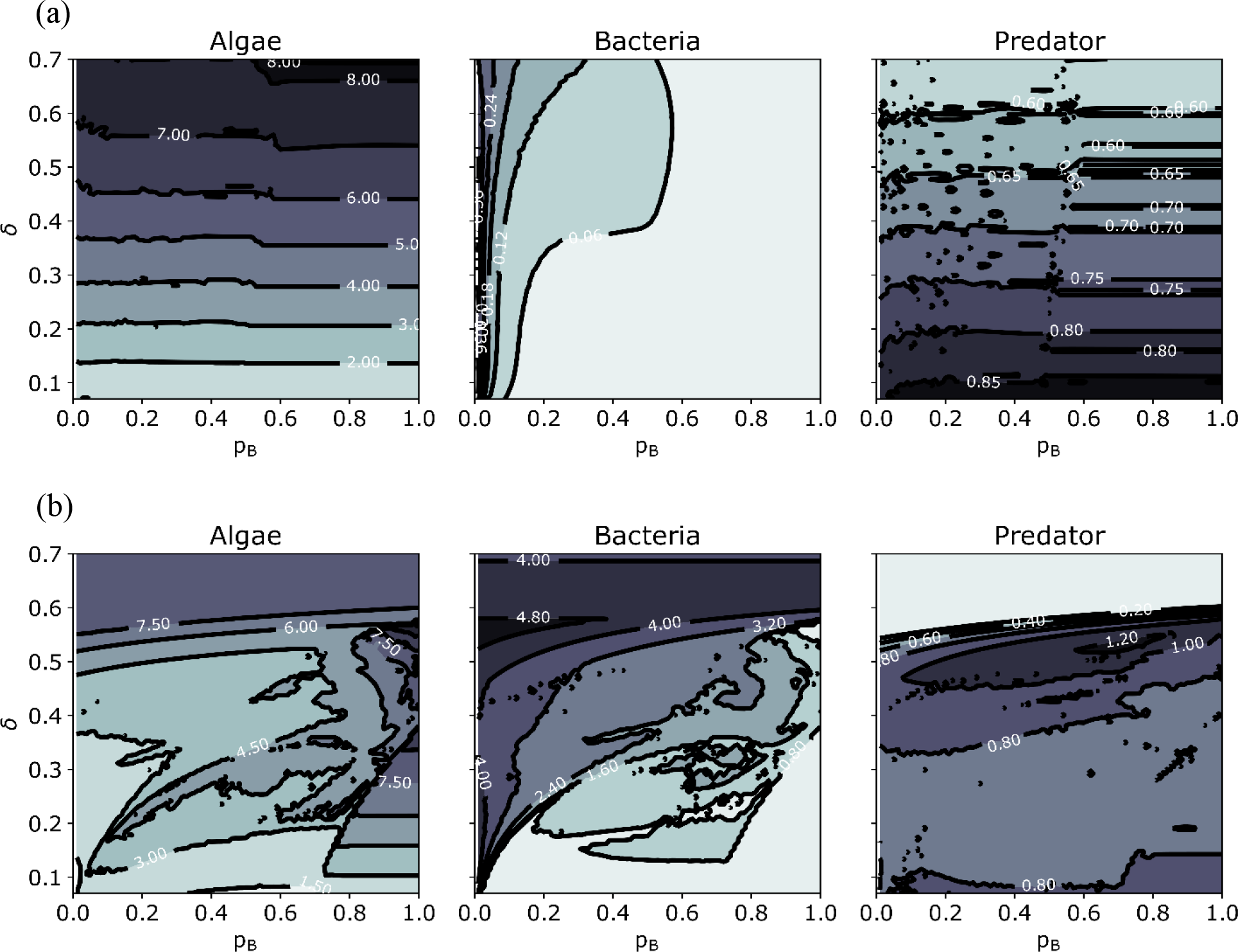
Mean biomasses for (a) suppressing conditions (algal exudation, predator excretion and bacterial growth efficiency are low) and (b) favouring conditions for bacteria (algal exudation, predator excretion and bacterial growth efficiency are high) for the parameter space spanned by the chemostat dilution rate δ and the degree of bacterivory of the predator *p*_*B*_. Colours correspond to different biomass levels [mgC/L] in the individual plots as the biomass ranges vary largely.

In contrast, under bacteria-favouring conditions bacteria reach considerable mean biomasses which are highest at low degrees of bacterivory and high dilution rates (Fig. 2b). An increasing degree of bacterivory results in lower bacterial and higher predator biomass. The algal biomass increases due to a release from competition. At very strong bacterivory and low dilution rate bacterial mean biomass becomes negligible in favour of the algae. The predator goes extinct if the dilution rate exceeds its maximal realized per capita growth rate. The dilution rates that the predator can withstand increase with stronger bacterivory.

The effect size of bacteria represented by the logarithmic ratio of mean biomasses in simulations with and without bacteria provides a direct measure of the impact of bacteria on mean biomasses (Fig. 3, Suppl. Fig. 3). While for suppressing conditions the bacterial biomass and thus the effect size of bacteria is negligible throughout the parameter space (Suppl. Fig. 3), interesting patterns emerge for favouring conditions (Fig. 3). Here, the total biomass (algae, rotifers and bacteria) decreases strongly if bacterivory and dilution rates are at intermediate levels, which originates from low algal biomasses that are not compensated by the bacteria and the biomass increase of the predator.

**Figure 3.**
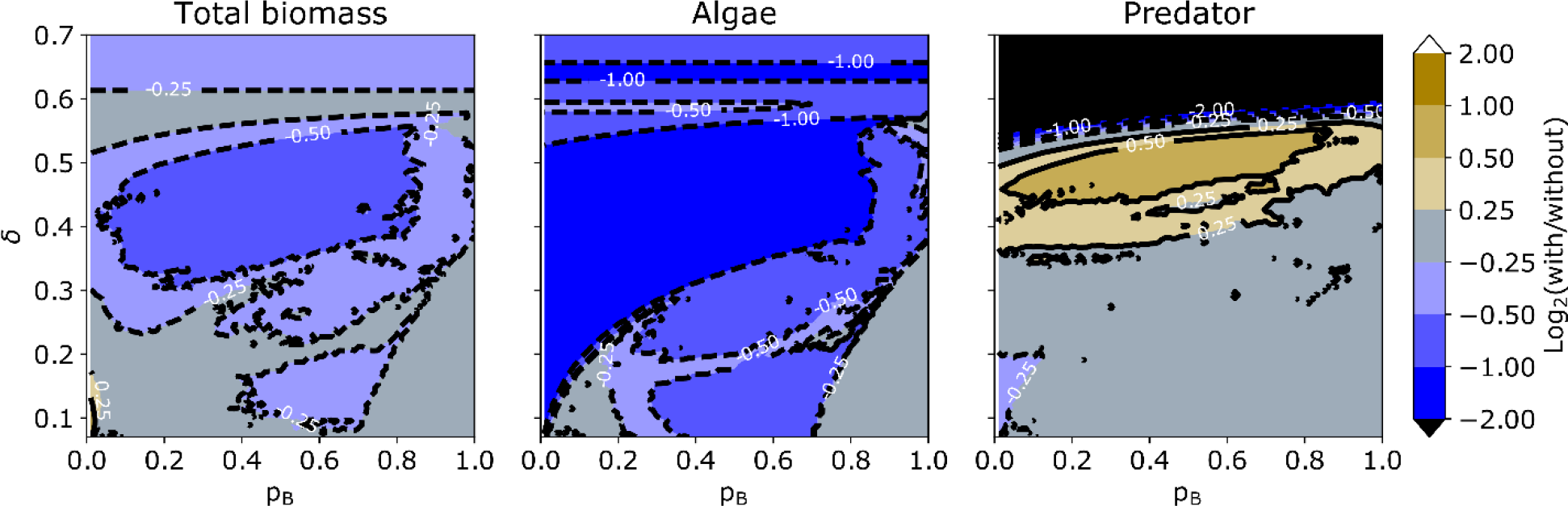
Effect size of bacterial presence under bacteria-favouring conditions. The effect size is defined as the logarithmic ratio to base 2 of the mean biomasses with and without bacteria. The presence of bacteria often decreases algal and total biomass but increases predator biomass. Under suppressing conditions the bacteria have only little effect (see Supplementary Figure 3).

### Effect of bacteria on population dynamics

Population dynamics are often used as fingerprints of biological interactions. To study how they are affected by the presence of bacteria, we scanned the parameter space of Fig. 2 and Fig. 3 for the type of population dynamics for the bacteria-favouring scenario where bacteria have a significant effect on the mean biomasses (Fig. 4). We found a complex pattern of alternating regions of regular and non-regular dynamics (Fig. 4a). At low dilution rates and high to intermediate degrees of bacterivory the population dynamics are fairly simple (panels i and ii in Fig. 4b). Within a cycle the algae establish first, nitrogen declines and organic carbon accumulates which allows the bacteria to increase as well. Finally, the predator reaches high biomasses by grazing down both algae and bacteria. This releases the nitrogen, the predator declines and the whole cycle starts again. However, if the degree of bacterivory is too high, the bacteria go extinct (as in panel i). These classic predator-prey cycles can easily become highly complex, driven by the interaction of direct and indirect competition between algae and bacteria (panels iii, iv, and v). For broad parameter ranges multiple maxima occur within one repetitive unit and partly the dynamics become irregular, i.e. no repetitive unit can be found in the time-series of the biomasses. At high dilution rates the cycle amplitude decreases (panel vi) and eventually the dynamics reach a steady state (panel vii). If the predator goes extinct, algae and bacteria continue to coexist in a steady state (panel viii).

**Figure 4.**
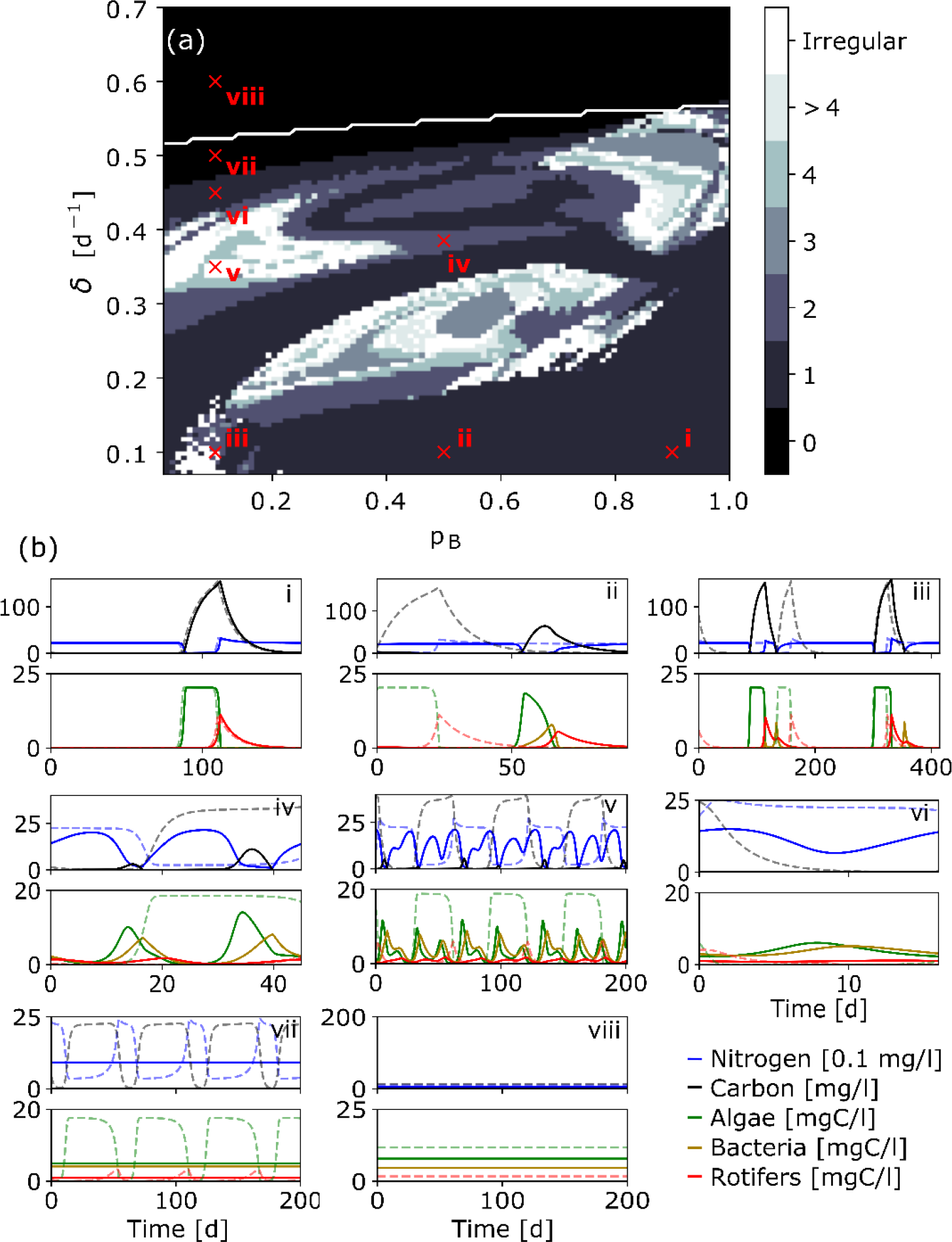
Population dynamics under favouring conditions. (a) Population dynamics as determined from the number of algal peaks within a repetitive unit. If no such unit was found the dynamics are classified as irregular. Mean predator biomass drops below 10^−30^ mgC/L above the white line. (b) Time series of one repetitive unit, if existent, for the marked parameter combinations. For steady states and irregular dynamics a fixed timespan of 200 days is plotted. Full and dashed lines correspond to runs with and without bacteria present, respectively. The dynamics can become highly complex, unless bacteria are grazed down by the predator.

Notably, for a fairly low degree of bacterivory and a low dilution rate, the rotifers are able to increase for a second time within one cycle even though the algae are already at a density too low to support the rotifers (panel iii). This second predator peak is mainly realized from grazing on bacteria.

### Explanation of results

The combined effect of dilution rate and degree of bacterivory can be understood by shifts in the balance between bottom-up and top-down control (Fig. 4). At high dilution rates and low degrees of bacterivory the predator is strongly limited in its net growth and the prey becomes more bottom-up limited. Thus, its cycle amplitudes decrease and mean biomasses increase. The low top-down control allows the prey to first deplete the resources before being washed out, in parts of the parameter space for multiple times during one predator cycle, before the predator has caught up and finally grazes down the prey. Within this first phase of low top-down control competition between algae and bacteria alternates with algae supporting bacterial growth through the release of organic carbon, which explains the complex multi-cycle patterns (Fig. 4). At low dilution rates and high degrees of bacterivory the top-down control increases as the predator is able to exert a considerable predation pressure on both algae and bacteria, thus forcing the system into more regular predator-prey cycles.

### Effect of nutrient inflow concentration

Similar to the above results, also the parameter space spanned by nutrient inflow concentration and degree of bacterivory is composed of regions of different bottom-up – top-down balances. As the inflow concentration increases the chemostat system is enriched and all mean biomasses increase (Suppl. Fig. 4a). An increasing degree of bacterivory has a similar effect for the predator as it broadens its food spectrum. Also, higher degrees of bacterivory suppress bacteria and favour algae in their apparent competition. Thus, if both parameters are low there is strong bottom-up control and the effect size of bacteria on the total biomass and the algae is small (Suppl. Fig. 4b). If nutrient inflow concentration and degree of bacterivory are high this results in a strong top-down control which again decreases the effect size of bacteria on the total biomass and the algae. At intermediate parameter ranges, however, both total biomass and algae are strongly negatively affected by bacteria. The effect size of bacteria on the predator behaves contrary. In the parameter regions of high bottom-up control and high top-down the predator is affected negatively, whereas it largely benefits from the bacteria in the intermediate region.

The dynamic pattern approximately reflects these three regions with simpler dynamics at strong forcing and more complex dynamics in the intermediate regime (Suppl. Fig. 4c).

## Discussion

Chemostat experiments, particularly with phytoplankton-zooplankton systems, are often employed to resolve ecological and evolutionary questions regarding predator-prey interactions. However, bacteria are omnipresent actors in nature. In this paper we argue that it may be indispensable to either include bacteria in the interpretation of study results or to take applicable measures to minimize their effect.

Using a simplified, static carbon-flux model as well as a mechanistic, dynamic chemostat model, which has been parametrized closely to typical experimental systems, we show that bacteria are able to strongly impact predator production, biomass levels and population dynamics in chemostat experiments. Under bacteria-suppressing conditions, i.e. if specific physiological properties of the organisms reduce the production and utilization of organic carbon, we expect bacteria to generally play only a minor role, if at all. Under bacteria-favouring conditions, however, predator production is substantially increased by the presence of bacteria. It is important to note that the contribution of bacteria to predator ingestion varies in time and thus temporally exceeds the mean values predicted by the static carbon-flux model. From the dynamic model we see that the effect of bacteria on the biomasses and particularly the population dynamics in the chemostat strongly depends on the experimental conditions, i.e. the dilution rate and nutrient inflow concentration, as well as the degree of bacterivory of the predators.

### Impact of the bacterial pathway on the food web structure

The shift of biomass from algae to bacteria at intermediate dilution rates and bacterivory decreases the total biomass in the chemostat when comparing systems with and without bacteria present. Here, the biomass of the bacteria and the biomass increase of the predator are not sufficient to compensate for the biomass losses of the algae as the bacterial pathway in the food web includes bacterial respiration as an additional loss-term along which biomass is irretrievably lost. This may obstruct predictions for biomass yield and energy balances in aquatic mass cultures if bacteria were not considered (Hino et al. 1997). On the other hand, this pathway increases the predator biomass as now algal exudates and predator excretion, which are lost without bacteria, are recycled by the bacteria and may be used by the predator, thus increasing the efficiency by which primary production is transferred to the predator.

### Importance of bacteria for population dynamics

While for high degrees of bacterivory and low dilution rates we observed regular predator-prey cycles, the dynamics can become highly complex for intermediate parameter regions. Within one pronounced and experimentally detectable cycle of the predator multiple cycles of algae and bacteria can occur. At low bacterivory and low dilution rates the overall dynamics resemble those without bacteria at first glance. The only indication of bacteria having an effect in this region is the second predator peak, which cannot be explained without considering bacteria in an experimental chemostat system and instead might lead to wrong conclusions.

Population time-series obtained by chemostat experiments occasionally are quite irregular (Bengfort et al. 2017). It remains to be studied whether this irregularity might be just a more complex attractor similar to the ones observed in this study. Bacteria could thus be an overlooked actor in chemostat experiments responsible for unexpected complexity of population dynamics.

### Implications for improvement of experimental design

Our study enables us to propose means for reducing the impact of bacteria in chemostat studies that explicitly focus on phytoplankton-zooplankton interactions by adjusting their design accordingly. Aside from the easily implemented measure to reduce the dilution rate, which enables a stronger response by the predator, also the ability to ingest bacteria should be taken into account when the predator species is selected. Instead of intuitively using species incapable of ingesting bacteria (e.g. numerous *rotifers*, Arndt 1993), predator species with a high degree of bacterivory could be the preferred choice (e.g. many *ciliates*, Sherr and Sherr 1987).

Bacteria can affect phytoplankton-zooplankton interactions in a chemostat by two mechanisms: (i) by competing for nutrients with the algae and (ii) by contributing to predator production. When we considered the effect sizes as a measure of the ratio of average biomasses with and without bacteria the zero-bacterivory limit corresponds to the case when only competition is at play. A high degree of bacterivory, however, includes the effect of both competition and predator divergence. Since the effect sizes do not vanish towards low degrees of bacterivory we see that bacteria have a considerable competitive impact on the algae and thus affect the food web even if they do not contribute to the production of the predator. A high degree of bacterivory of the predator minimizes the competitive impact of bacteria and thus decreases the effect of bacteria in chemostat experiments.

Here we argue that bacteria are an unavoidable and inherent actor in phytoplankton-zooplankton chemostats, whose impact may be minimized by choosing the right experimental setup. Thereby we should keep in mind that – up to now overlooked – bacteria might have some impacts on population dynamics and species coexistence that are comparable to the previously overlooked effects of rapid evolution (Yoshida et al. 2003).

Our study shows that only with an appropriate choice of the predator species and an appreciation for the presence and role of all important actors we can correctly interpret phytoplankton-zooplankton chemostats and use them to study complex predator-prey interactions.

## Acknowledgements

This work was funded by DFG (WA 2445/8-1, WA 2445/11-1, GA 401/26-1) as part of the Priority Programme 1704 (DynaTrait).

## Appendix 1 Static model

**Supplementary Figure 1.**
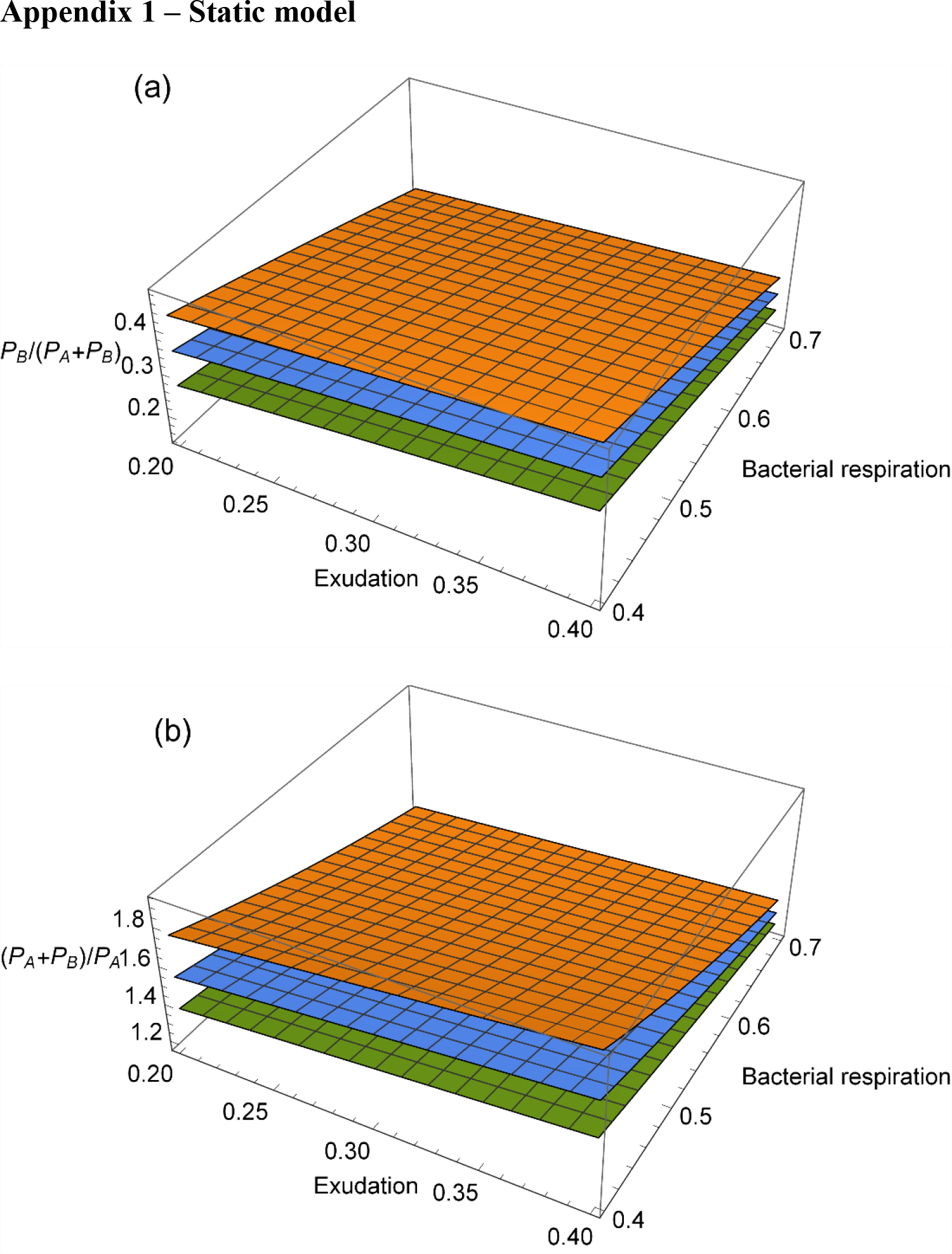
The behaviour of the static model for the three physiological parameters that determine the exudation, i.e. the fraction of carbon that is exudated by algae *e*, the bacterial respiration, i.e. the fraction of carbon taken up by bacteria that is respired *r*_*B*_ and the assimilation efficiency of predators ɛ. The assimilation efficiency is set to 0.4 (orange, top), 0.55 (blue, middle) and 0.7 (green, bottom). The respiration of the predator is set to r_*R*_ = 0.5. (a) Predator production derived from bacteria (*P*_*B*_) relative to total production. (b) Total predator production with bacteria present in the chemostat relative to without bacteria.

## Appendix 2 Derivation of the exudation

We assume that the per capita rate *r*_*growth*_ at which an alga grows in units of carbon is determined by a three-step process. First, organic carbon has to be fixed via photosynthesis which happens at rate *r*_*fix*_. Secondly a portion of this organic carbon is exudated at rate *e*_*dyn*_ *r*_*fix*_. Finally, the remaining carbon (1 − *e*_*dyn*_) *r*_*fix*_ may be used to build new biomass, depending on the nitrogen availability given by the Monod term 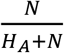.

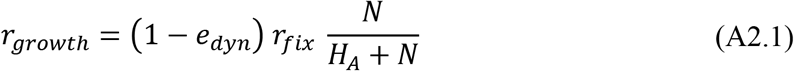

The portion of organic carbon that is exudated increases under nitrogen limitation, given by 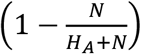. We assume that the exudation *e*_*dyn*_ is a linear function of the nitrogen limitation and bounded between a minimum *e*_*min*_ and a maximum *e*_*max*_ (Supplementary Figure 2a).

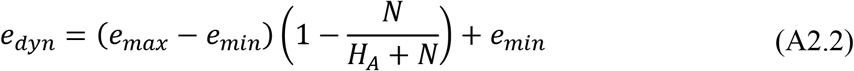

The flux of exudated carbon equals

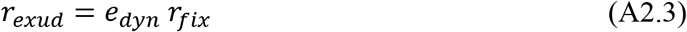

We assume that the production of organic carbon operates at a fixed rate. It is measured if nitrogen is not limiting, i.e. 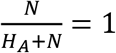, as the maximum per capita growth rate in units of carbon and it follows from Eqs. A2.1 and A2.2 that

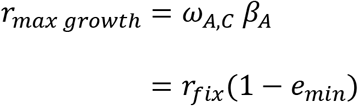

and therefore

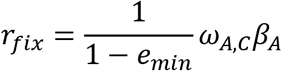

With Eq. A2.3 the per capita exudation rate in units of carbon becomes

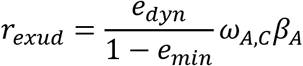

The per capita growth rate under nitrogen limitation with exudation included (Eq. A2.1) thus becomes

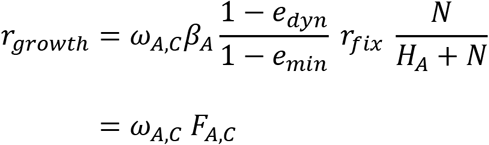

For plots of the exudation rate and growth rate at different maximal exudation ratios see Suppl. Fig. 2.

**Supplementary Figure 2.**
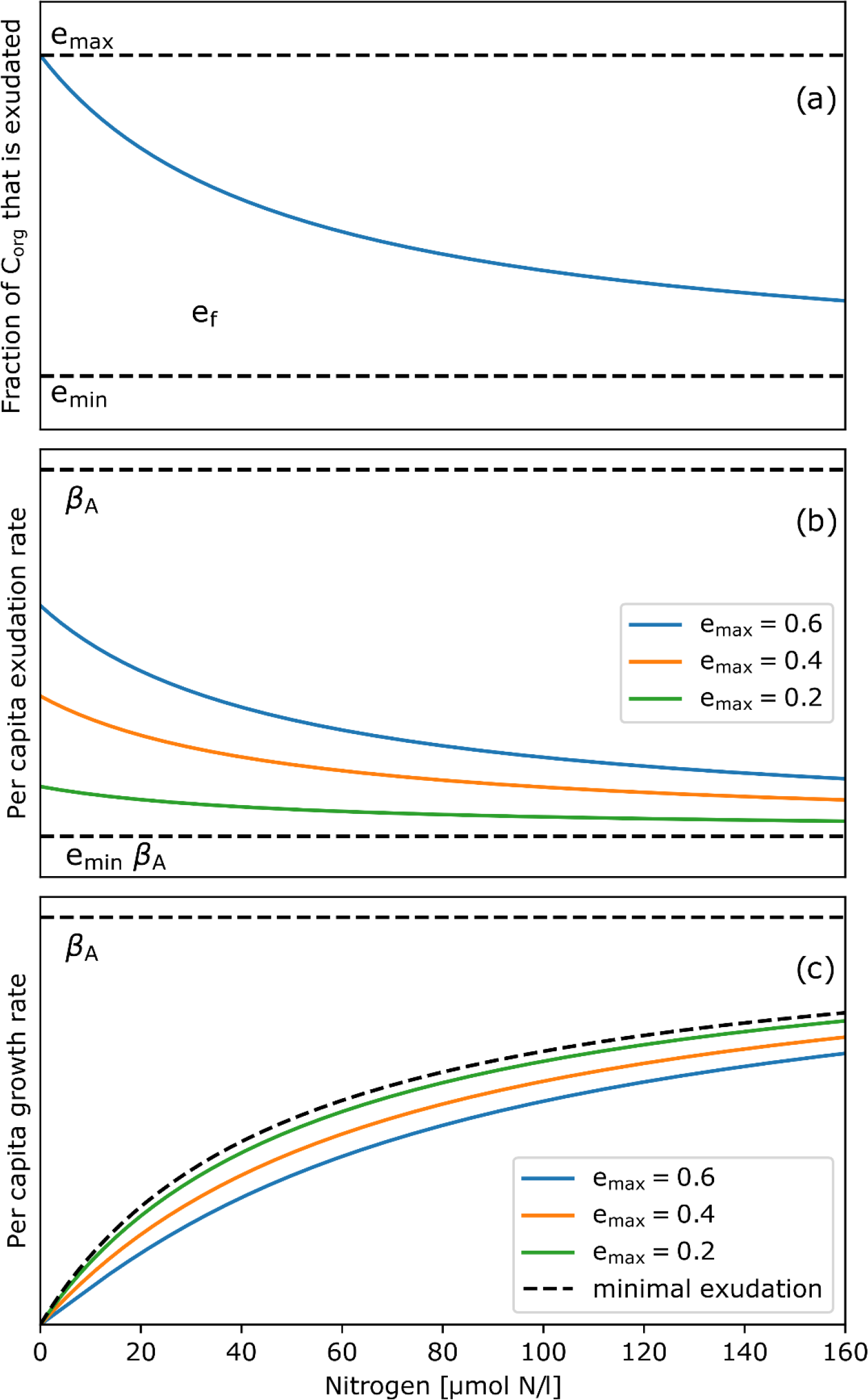
Effect of the exudation model for a half-saturation constant of *H*_*A*_ = 49μmolN/L and a minimal exudation of *e*_*min*_ = 0.1. (a) Fraction of fixed carbon that is exudated. (b) Carbon exudation rate relative to the realized per-capita growth rate β_A_. (c) Per capita growth rate. The dashed black line represents growth that is only affected by minimal exudation.

## Appendix 3 Effect size for suppressing conditions

**Supplementary Figure 3.**
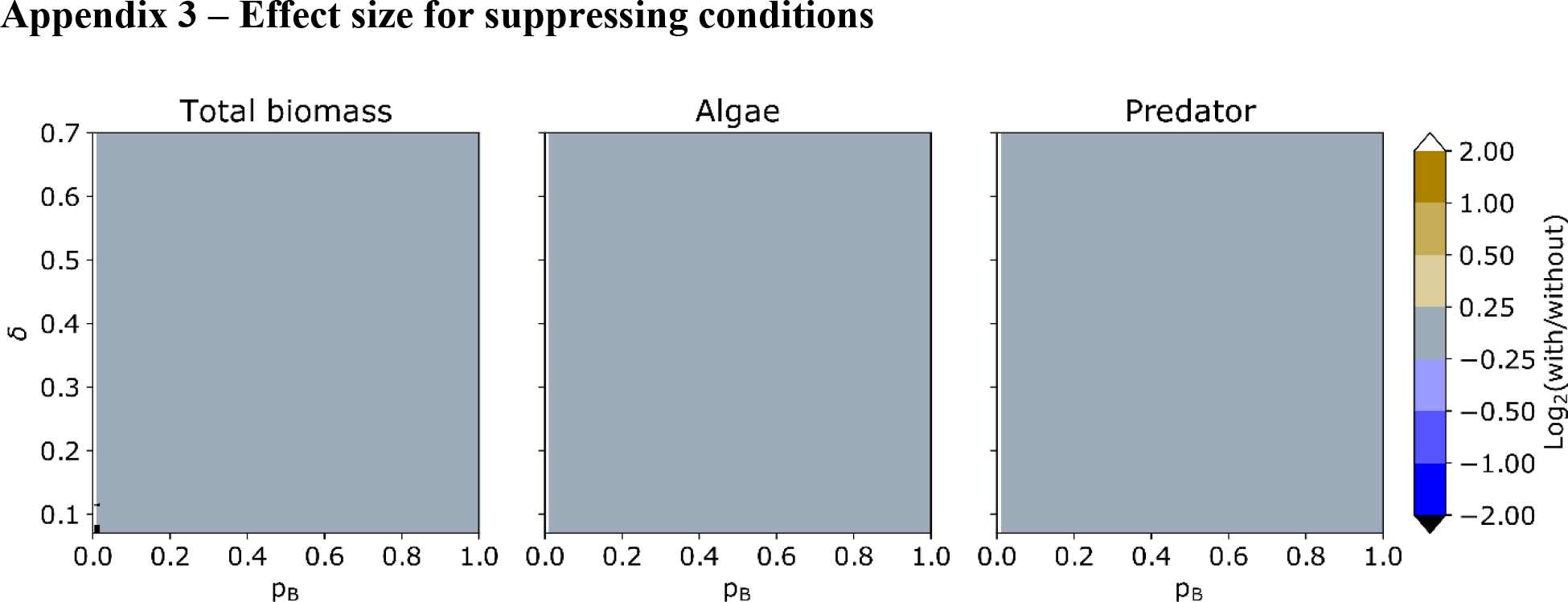
Effect size of bacterial presence under bacteria-suppressing conditions: *e*_*max*_ = 0.2, *r*_*B*_ = 0.7 and ɛ = 0.7. The effect size is defined as the logarithmic ratio to base 2 of the mean biomasses with and without bacteria.

## Appendix 4 Effect of nutrient inflow concentration

**Supplementary Figure 4.**
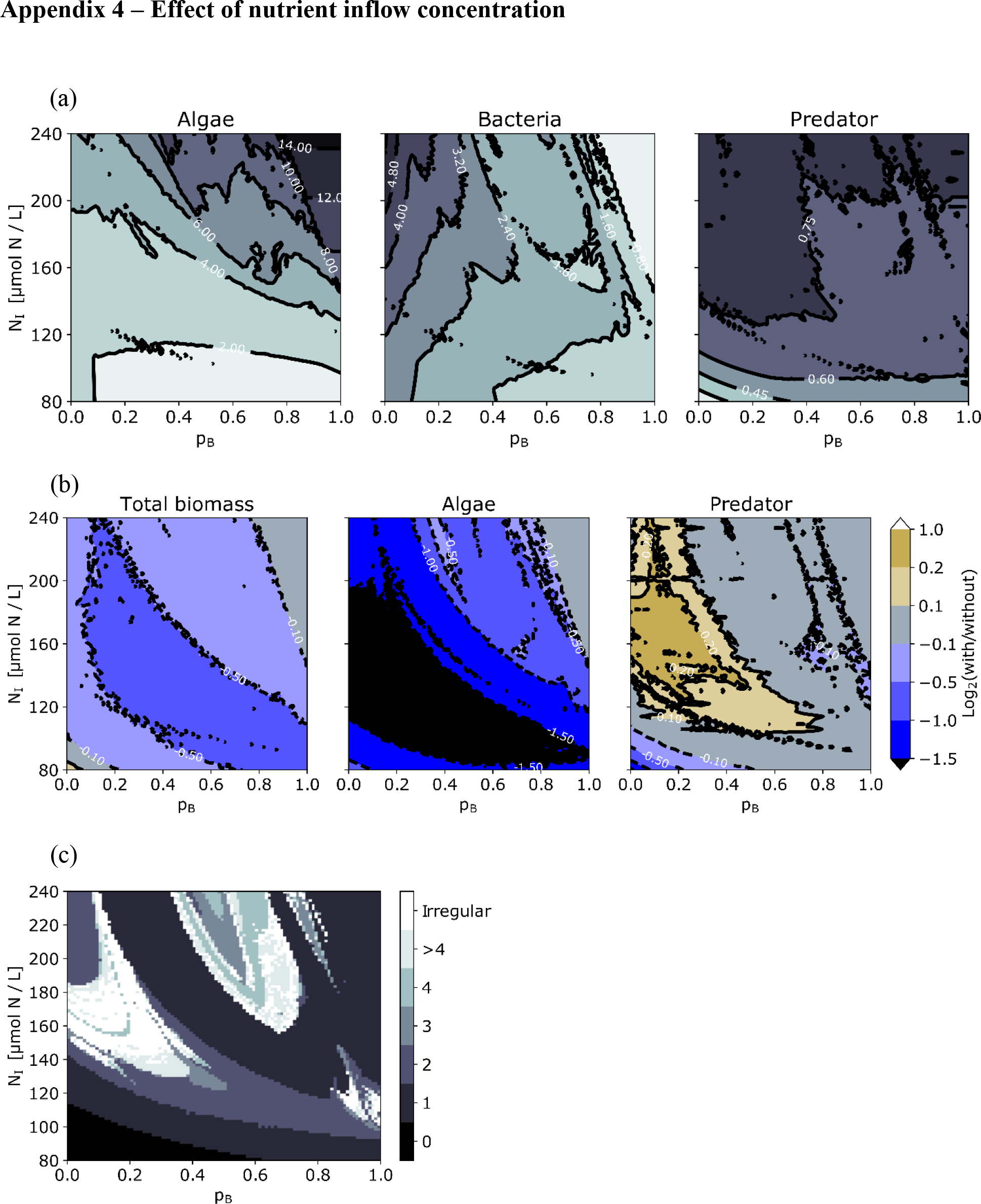
Parameter space spanned by degree of bacterivory and nutrient inflow concentration under favouring conditions at a dilution rate of *δ* = 0.35 *d*^−1^ for (a) mean biomasses, (b) effect size of bacterial presence and (c) types of dynamics characterized by the number of algae maxima per repetitive unit.

